# Microtubule-dependent transport of arenavirus matrix protein demonstrated using live-cell imaging microscopy

**DOI:** 10.1101/665521

**Authors:** Yuki Takamatsu, Junichi Kajikawa, Yukiko Muramoto, Masahiro Nakano, Takeshi Noda

**Affiliations:** Laboratory of Ultrastructural Virology, Institute for Frontier Life and Medical Sciences, Kyoto University, Japan; Institut für Virologie, Philipps-Universität Marburg, Germany

## Abstract

Lassa virus (LASV), belonging to the family *Arenaviridae*, causes severe haemorrhagic manifestations and is associated with a high mortality rate in humans. Thus, it is classified as a biosafety level (BSL)-4 agent. Since counter measures for LASV diseases are yet to be developed, it is important to elucidate the molecular mechanisms underlying the life cycle of the virus, including its viral and host cellular protein interactions. These underlying molecular mechanisms may serve as the key for developing novel therapeutic options. Lymphocytic choriomeningitis virus (LCMV), a close relative of LASV, is usually asymptomatic and is categorised as a BSL-2 agent. In the present study, we visualised the transport of viral matrix Z protein in LCMV-infected cells using live-cell imaging microscopy. We demonstrated that the transport of Z protein is mediated by polymerised microtubules. Interestingly, the transport of LASV Z protein showed characteristics similar to those of Z protein in LCMV-infected cells. The live-cell imaging system using LCMV provides an attractive surrogate measure for studying arenavirus matrix protein transport in BSL-2 laboratories. In addition, it could be also utilised to analyse the interactions between viral matrix proteins and the cellular cytoskeleton, as well as to evaluate the antiviral compounds that target the transport of viral matrix proteins.

## Introduction

The family arenavirus contains at least 41 recognised species with single-stranded ambisense RNA genomes (1). Their considerable impact on human health should not to be underestimated, as described in a review by Wolff, *et al.* (2). In this family, the Lassa virus (LASV), which causes the haemorrhagic Lassa fever, is a major public health concern due to its potential to cause international epidemics (3–8). LASV is prevalent in West Africa, and over 500 cases of Lassa fever were confirmed in Nigeria in 2018 (8). The only available drug for the treatment of Lassa fever is ribavirin, which despite being partially effective, has significant side effects, including haemolytic anaemia (9). Due to the lack of effective prevention and counter measures, LASV is classified as a biosafety level 4 (BSL-4) microorganism. A BSL-4 facility requires specific biosafety management and laboratory design, including controlled access and air and drainage systems, which must meet international and national regulations (10). Lymphocytic choriomeningitis virus (LCMV), a close relative of LASV, is generally asymptomatic in healthy individuals and is categorised as a BSL-2 microorganism. Thus, LCMV can be used as an alternative to analyse the molecular mechanisms of each replication step during LASV infection (11–13).

Arenavirus particles have a spherical or pleomorphic morphology, with a diameter ranging from 60 to 300 nm. Two segments (small and large) of viral RNA encode the following four viral proteins: the surface glycoprotein GP and nucleoprotein NP are encoded by the small segment, while the large segment encodes the polymerase L and matrix protein Z (Fig. 1a). The ribonucleoprotein complexes (RNPs) are formed by genomic RNA segments associated with NP and L (Fig. 1b). Transcription and replication of viral genome occurs in the host cell cytoplasm, and are achieved by the RNPs (14). Z protein downregulates these processes through its interaction with the viral polymerase (13, 15, 16). In addition, Z protein plays a crucial role in virion assembly and release (2, 13, 17). Although the importance of viral late domain motifs in Z protein and the interaction of Z protein with the cellular endosomal sorting complexes required for transport (ESCRT) for virion formation and budding have been well studied (2, 13, 17-22), the transport of Z proteins is been yet fully understood. In the present study, we performed live-cell imaging analyses of Z protein transport in LCMV-infected cells and characterised the movement both over short time period (within 3 min, denoted “short duration”) and long time period (up to 12 h, denoted “long duration”). We also characterised the movement of LASV Z protein in plasmid-transfected cells.

**Figure 1.**
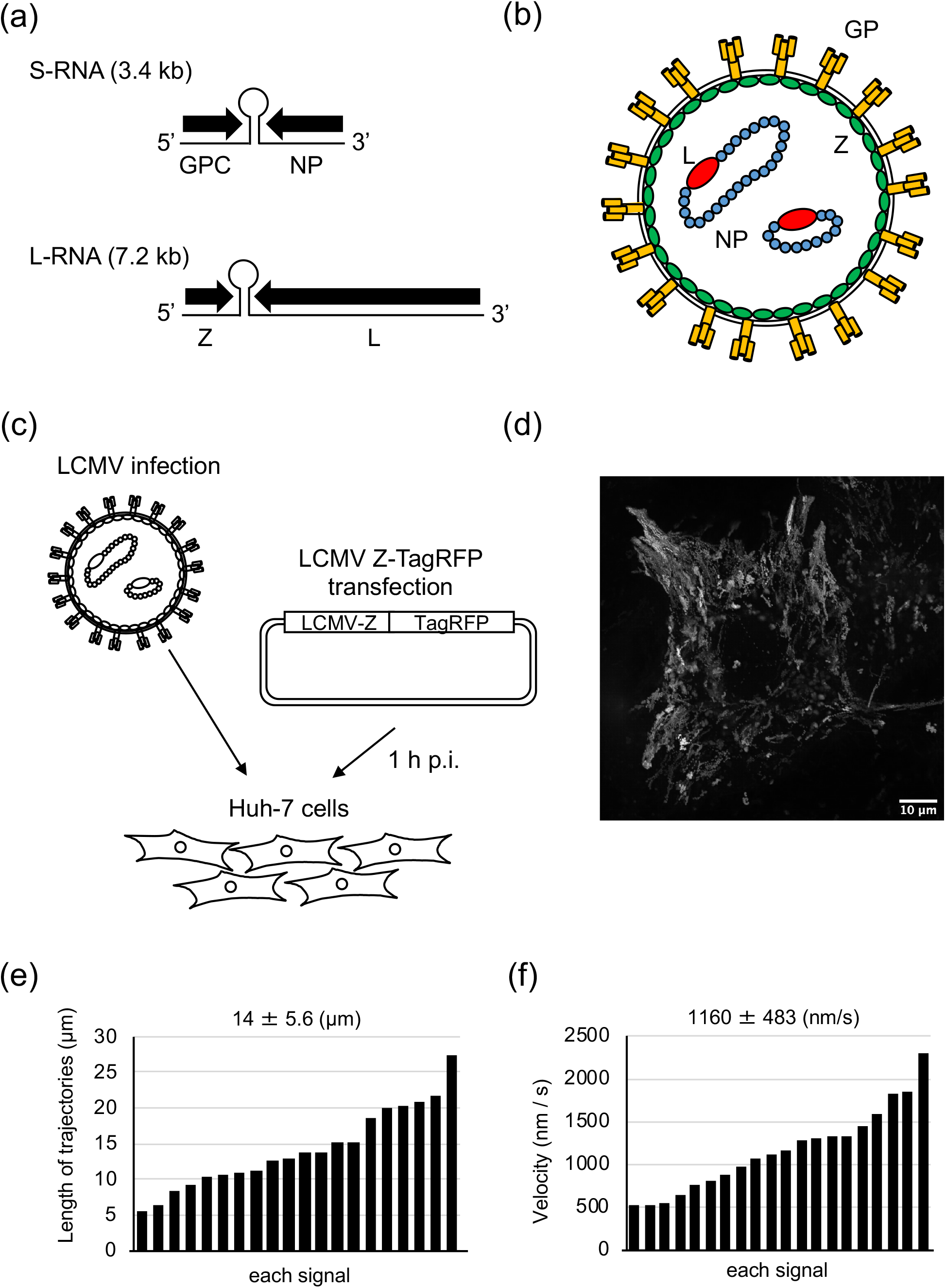
Schematic diagram of arenavirus genome, virus particle, and live-cell imaging for Z proteins in Lymphocytic choriomeningitis virus infection. (a) Genome organization of arenaviruses. Arenaviruses genome composed of the small (S) RNA segment and the large (L) RNA segment, utilizing an ambisense coding strategy. The open reading frames are separated by intergenic regions. The S segment encodes NP and GPC. The L segment encodes L and Z. (b) Structure of arenavirus particle. RNPs are formed with viral RNA encapsidated by multiple NPs, and are associated with a polymerase L. Z protein forms a matrix layer underneath the viral membrane, where GPs are embedded. (c) Experimental setting for detecting Z protein transport. Huh-7 cells were infected with LCMV at MOI 1 and transfected with a plasmid encoding LCMV Z-TagRFP at 1 h p.i. Live-cell imaging microscopy was conducted 24 h p.t. (d) LCMV Z protein intracellular movement. The picture shows the maximum-intensity projection of time-lapse images of cells recorded for 3 min; images were captured every 1 s. (e) The length of selected Z proteins trajectories is shown (n=20). The number indicates mean ± SD (μm). (e) The velocity of selected RNPs is shown (n=20). The number indicates mean ± SD (nm/s).

## Materials and methods

### Cell culture

Huh-7 (human hepatoma) cells were maintained in Dulbecco’s Modified Eagle Medium (DMEM, Life Technologies) containing 10% (vol/vol) fetal bovine serum (FBS, PAN Biotech), 5 mM L-glutamine (Q; Life Technologies), 50 U/mL penicillin, and 50 μg/mL streptomycin (PS; Life Technologies), grown at 37 °C with 5% CO_2_. A maintenance medium, containing DMEM supplemented with 3% FBS and Q/PS, was used for cultivation after cell growth.

### Virus infection

LCMV Armstrong 53b strain (GenBank accession number: P18541) was propagated in BHK-21 cells. The titre was determined by a plaque assay using Vero cells (12). For live-cell imaging, 2 × 10^5^ Huh-7 cells were grown in a 35 mm dish (ibidi) with growth medium. Prior to virus inoculation, the growth medium was replaced with the maintenance medium (DMEM with Q/PS) and virus infection was achieved at a multiplicity of infection (MOI) of 1. After incubating for 1 h at 37 °C with 5% CO_2_, the virus inoculum was replaced with the maintenance medium.

### Plasmids and transfection

To generate the fluorescence-conjugated LCMV Z protein encoding plasmid (pCAGGS-Z-TagRFP), the TagRFP ORF was cloned in-frame to the 3’ end of the Z gene, as previously described (23). The fluorescence-conjugated LASV Z protein, pCAGGS-Z-GFP, was kindly provided by Dr. Kawaoka (Tokyo University). The plasmid DNA and transfection reagent TranSIT (Mirus) were mixed in 100 μL Opti-MEM without phenol red (Life Technologies) and added to the Huh-7 cells according to the manufacturer’s instructions.

### Live-cell imaging microscopy

Huh-7 cells at a cell density of 2 × 10^5^ were seeded in a 35 mm dish (ibidi) and cultivated in DMEM/PS/Q with 10% FBS. To analyse the movement of LCMV Z protein, first the cells were infected with the virus, as described above. At 1 h post-infection (h.p.i.), pCAGGS-Z-TagRFP (1 μg) was transfected. The medium was removed at 24 h post-transfection (h.p.t.) and 500 μL of live-cell imaging medium (containing CO_2_-independent Leibovitz’s medium (Life Technologies) with PS/Q, non-essential amino acid solution, and 0.5 % (vol/vol) FBS) was added. To analyse the movement of LASV Z proteins, pCAGGS-Z-GFP (1 μg) was transfected. The same procedures as those described above were carried out, except for viral infection. The live-cell time-lapse experiments were recorded using GE healthcare Delta Vision Elite with a 60× oil objective in a biosafety level-2 laboratory. The movement of the proteins for the short duration was recorded at 1 s intervals, for a total of 2-3 min. The movement of the protein for the long duration was recorded at 15 min intervals, for a total of 12 h.

### Treatment of cells with cytoskeleton-modulating drugs

Cells were treated with 15 μM nocodazole (Sigma), 0.3 μM cytochalasin D (Sigma), or 0.15% dimethyl sulfoxide (DMSO, Sigma) (24). These chemicals were added to the live-cell imaging medium 3 h prior to observation.

### Image processing and analysis

The images and movie sequences obtained through the observation were processed and analysed manually using the Fiji plug-in “MTrackJ” (25, 26).

## Results and Discussion

### Visualization of Z protein transport in LCMV-infected cells

Although the transport of LASV Z protein is known to be regulated by a microtubule-dependent motor protein, KIF13A (27), the characteristics of arenavirus Z protein transport during viral infection have not yet been fully elucidated. In cells infected with the Ebola and Marburg viruses, fluorescence-conjugated matrix proteins have been previously used to visualise the transport of viral proteins in virus-infected cells (23, 24). In the present study, we used these live-cell imaging methods, previously used with filoviruses, to observe arenaviruses.

Huh-7 cells are flat in shape and are suitable for tracking the movement of proteins using live-cell imaging. First, these cells were infected with LCMV at a MOI of 1 and transfected with 1 μg of pCAGGS-Z-TagRFP at 1 h p.i (Fig. 1c). Next, time-lapse images were acquired at 1 s intervals for a total of 2-3 min, from 24 h p.t. The length of the trajectories for the short duration (2 min) ranged from 5 to 30 µm, with a mean length of 14 ± 5.6 μm (Fig. 1e). The velocity of transport was 500 to 2500 nm/s, with a mean speed of 1160 ± 483 nm/s (Fig. 1f), suggesting that the transport of Z proteins was likely mediated by microtubules rather than actin cytoskeletons.

### LCMV Z protein transport along polymerised Tubulin

Many viral proteins use the host cell cytoskeleton as a mode of transport (23, 28-30). Thus, cytoskeleton-modulating drugs are useful to analyse the intracytoplasmic transport of viral matrix proteins (24). Huh-7 cells were infected and transfected as described in Fig. 1c. The culture medium was then replaced at 24 h p.t. with a volume of 500 μL CO_2_-independent Leibovitz’s medium containing either 0.15% DMSO (control), 0.3 µM of cytochalasin D (actin depolymerizing drug), or 0.15 M of nocodazole (microtubule depolymerizing drug), and the cells were incubated for 3 h with each drug. Time-lapse images were subsequently acquired at 1 s intervals for a total of 2-3 min (Fig. 2a). Cytochalasin D-treated cells were found to not alter the characteristics of Z protein movement in comparison to the control cells (Fig. 2b and 2c, Supplementary Movie 2 and 3). The mean length of the trajectories was 15 ± 5.5 μm and 13 ± 6.3 μm for DMSO- and cytochalasin D-treated cells, respectively (Fig. 2e and 2f). The mean velocity of transport was 1130 ± 404 nm/s and 1101 ± 417 nm/s for for DMSO- and cytochalasin D-treated cells, respectively (Fig. 2h and 2i). Nocodazole treatment was found to immediately induce the cessation of Z protein movement, without any long distance transport (Fig. 2d, 2g, 2j, Supplementary Movie 4). These results indicate that arenaviruses Z proteins utilise polymerised microtubules for transport. The velocity of intracellular transport of LCMV Z proteins was within a similar range as intracellular-enveloped vaccinia virus transport, which is also mediated by microtubules (31–33).

**Figure 2.**
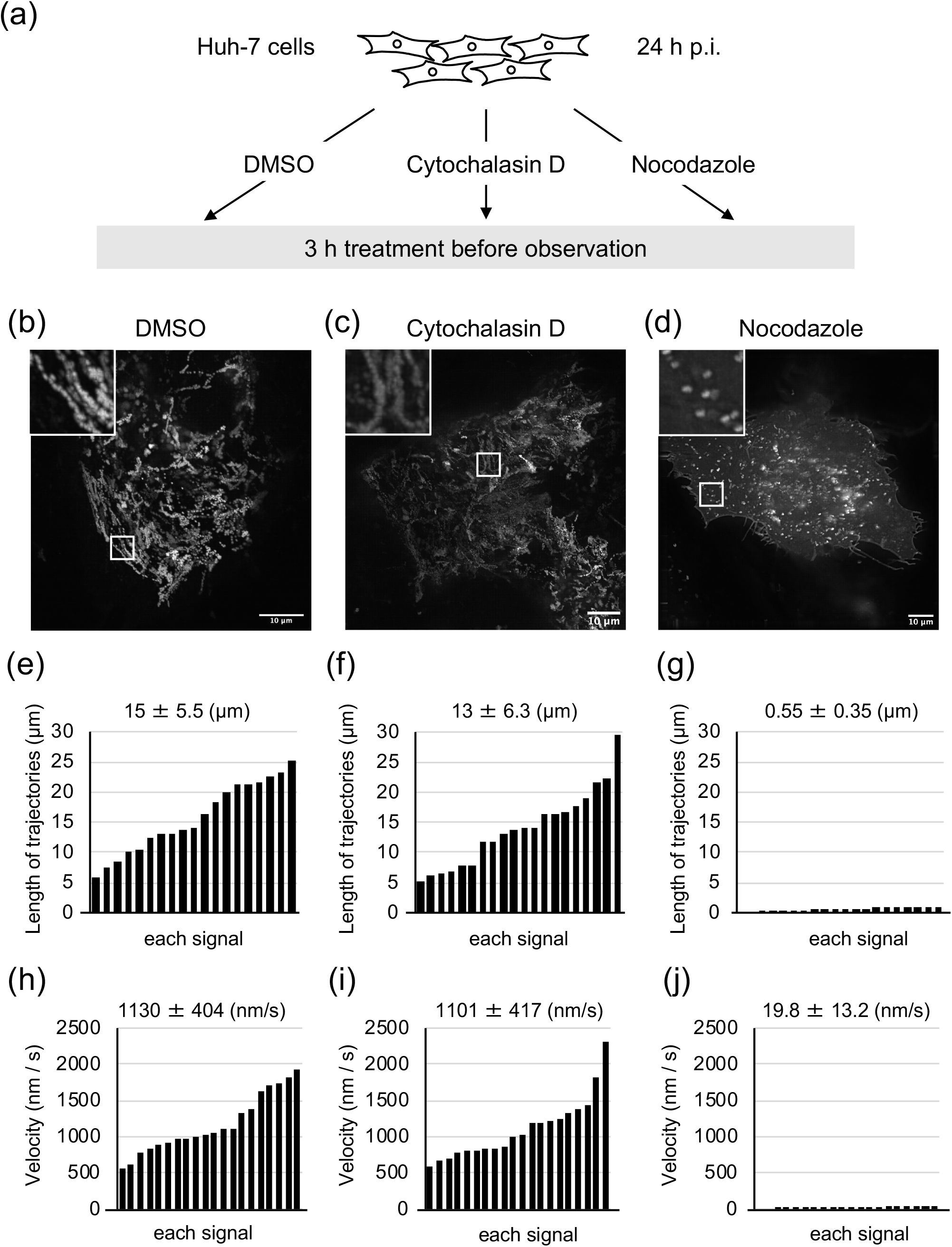
Characteristics of LCMV Z proteins transport in short duration. (a) Experimental setting to observe effects of cytoskeleton-modulating drugs on Z protein transport in LCMV-infected cells. Huh-7 cells were infected and transfected as described in Fig. 1(b), and treated either with DMSO (control), cytochalasin D, or nocodazole at 24 h p.i. After 3 h of treatment, observation was started. (b-d) Time-lapse images acquired in each drug-treated cell (b: DMSO, c: cytochalasin D, d: nocodazole). The pictures show the maximum-intensity projection of time-lapse images of cells recorded for 3 min; images were captured every 1 sec. (e-g) The length of selected Z proteins trajectories is shown in each drug treated cells (n=20, e: DMSO, f: cytochalasin D, g: nocodazole). The number indicates mean ± SD (μm). (h-j) The velocity of selected Z proteins transport is shown in each drug treated cell (n=20, h: DMSO, i: cytochalasin D, j: nocodazole). The number indicates mean ± SD (nm/s).

### LCMV Z protein transport over long duration

Next, we analysed Z protein transport over long duration in Huh-7 cells. Time-lapse images were acquired over a period of 12 h at 15 min intervals after treating the cells with cytoskeleton-modulating drugs for 3 h. Consistent with the short duration experiments, Z protein transport was not restricted in control-treated cells, but was strictly regulated in nocodazole-treated cells (Fig. 3a, b). These results indicate that microtubule polymerisation is the key modulator for Z proteins over long duration in LCMV-infected cells. These results correlated with previous findings, which showed that nocodazole treatment induced a significant reduction in arenavirus production (27).

**Figure 3.**
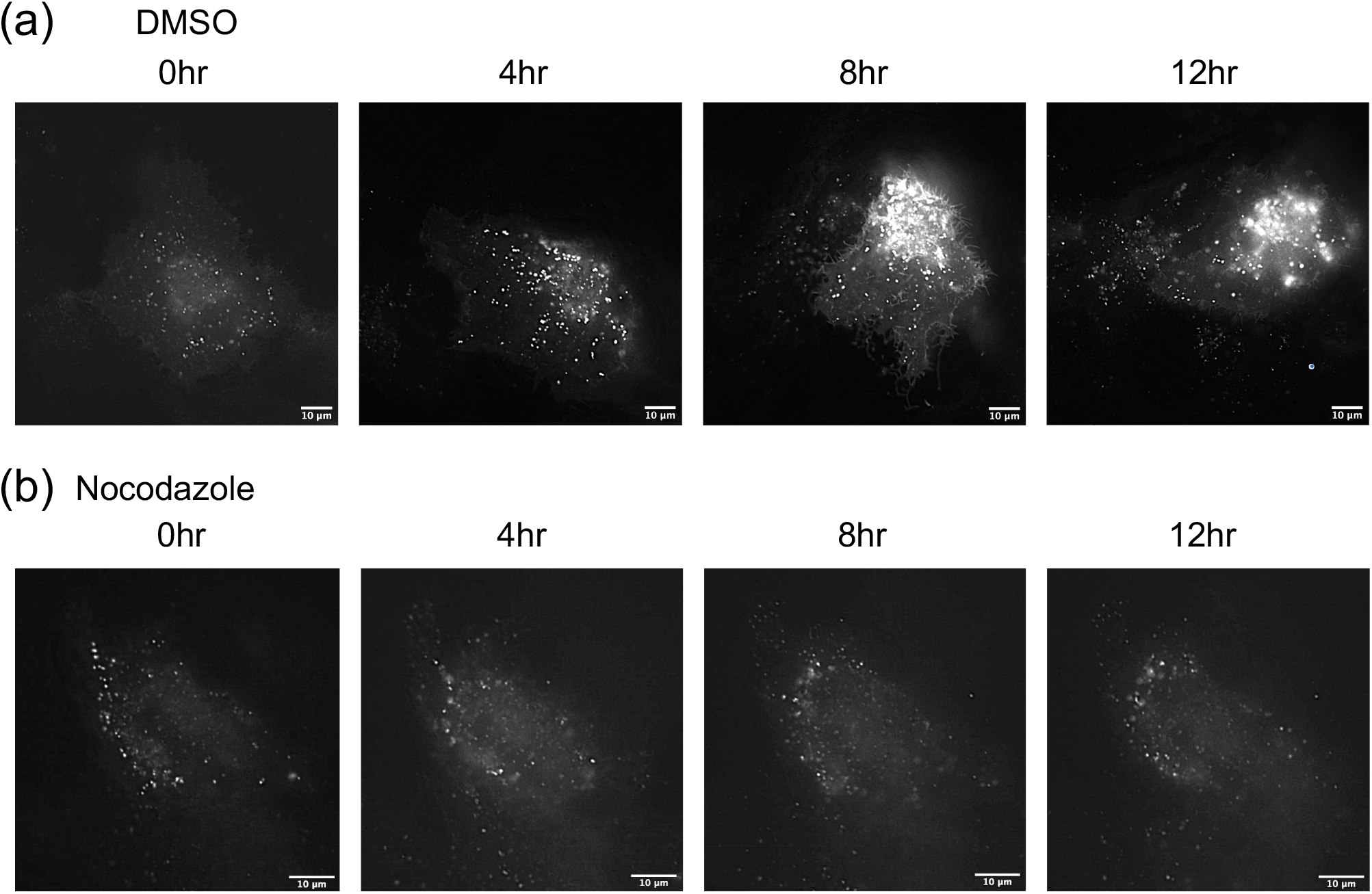
Z proteins transport in LCMV-infected cells over long duration. The effect of cytoskeleton-modulating drugs on LCMV Z protein transport for long duration. Huh-7 cells were infected and transfected as described in Fig. 1(b), and treated either with DMSO (a), or nocodazole (b) at 24 h p.i. The cells were observed for 3-15 h after drug treatment; images were captured every 15 min over a duration of 12 h. The pictures show the cells at the indicated time post-observation.

### Characterization of LASV Z protein transport in short and long durations

To evaluate the relevance of our assay for highly pathogenic LASV, we analysed the intracellular movement of Z protein in LASV Z protein-expressing cells (Fig. 4a). Huh-7 cells were first transfected with 1 μg of pCAGGS-Z-GFP, followed by cytoskeleton-modulating drugs, as described above. Although neither DMSO nor cytochalasin D influenced Z protein movement, treatment with nocodazole abolished Z protein movement in the short duration (Fig. 4b, c, d). The transport of Z proteins was found to be mediated via polymerised microtubules in the long duration (Fig, 4e, f). Thus, our analysis of Z protein transport during LCMV infection could serve as an effective surrogate method for analysing LASV Z protein transport during LASV infection in BSL-4 laboratories.

**Figure 4.**
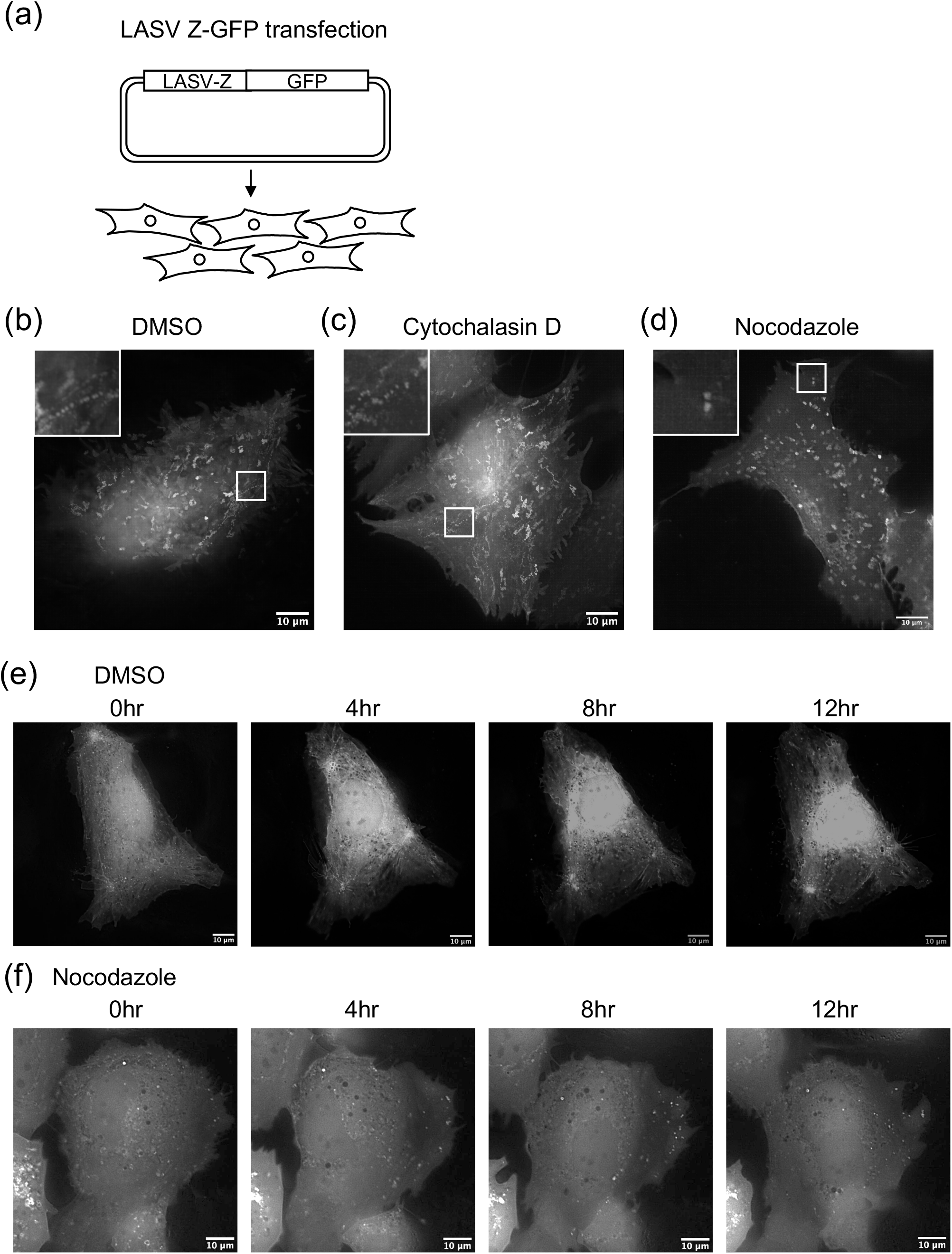
Characterization of Lassa virus Z proteins transport. (a) Experimental setting to observe LASV Z protein transport. (b-d) Huh-7 cells were transfected with pCAGGS-Z-GFP and treated either with DMSO (control), cytochalasin D, or nocodazole at 24 h p.i. Time-lapse images were acquired in each drug-treated cell (b: DMSO, c: cytochalasin D, d: nocodazole). The pictures indicate time-lapse images of cells recorded for 3 min; images were captured every 1 sec. (e,f) Effects of cytoskeleton-modulating drugs upon LASV Z proteins transport for long duration. Huh-7 cells were transfected as described in Fig. 4 (a), and treated either with DMSO (a), or nocodazole (b) at 24 h p.i. to the cells were observed for 3-15 h after drug treatment; images were captured every 15 min over a duration of 12 h. The pictures show the cells at the indicated time post-observation.

### Concluding remarks

Viruses utilise the cytoskeletons of infected cells to transport their components from the site of protein synthesis to the site of virus morphogenesis. In the present study, we established a live-cell imaging system for arenavirus Z protein transport using fluorescence-conjugated Z proteins. As reported previously, the trafficking of LASV Z proteins is regulated by a microtubule-dependent motor protein (27). Our results correlated with these findings and provided direct evidence that the transport of LCMV and LASV Z proteins is restricted by microtubule polymerization. The live-cell imaging approach developed here provides an effective method for analysing interactions between arenavirus Z proteins and cellular cytoskeletons, as well as for evaluating antiviral compounds that target the transport of arenavirus matrix proteins.

## Acknowledgments

The authors would like to extend their gratitude to Yoshihiro Kawaoka (Tokyo University) for providing the materials, and to Thomas Strecker (Philipps University Marburg, Germany) and Shuzo Urata (Nagasaki University, Japan) for the fruitful discussions.

## Funding

The work was supported by the Japan Society for the Promotion of Science JSPS (grant no. 18J01631 and 19K16666) (to Y.T.); by the AMED Research Program on Emerging and Re-emerging Infectious Diseases; by the AMED Japanese Initiative for Progress of Research on Infectious Disease for global Epidemic; by the JSPS Core-to-Core Program A, the Advanced Research Networks; by the Grant for Joint Research Project of the Institute of Medical Science, University of Tokyo; by the Joint Usage/Research Center Program of the Institute for Frontier Life and Medical Sciences Kyoto University; by the Daiichi Sankyo Foundation of Life Science, and by the Takeda Science Foundation (to T.N.).

